# Do more food choices lead to bad decisions? A case study in predacious ladybird beetle, *Propylea dissecta*

**DOI:** 10.1101/2023.11.03.565598

**Authors:** Lata Verma, Geetanjali Mishra, Omkar

**Affiliations:** Research Scholar, Ladybird Research Laboratory, Department of Zoology, University of Lucknow, Lucknow – 226007, India; Professor, Ladybird Research Laboratory, Department of Zoology, University of Lucknow, Lucknow – 226007, India

**Keywords:** Food preference, pollen grains, dietary regime, adult food choice

## Abstract

Understanding why animals choose one food over another is one of the key questions underlying the fields of behaviour ecology. This study aims to test if ladybird beetles, *Propylea dissecta* Mulsant (Coleoptera: Coccinellidae) can forage selectively for nutrients in order to redress specific nutritional imbalances to maximise their fitness. The general approach was, first, to manipulate the nutritional status of the predator by rearing them in five separate pre-treatment dietary groups from first instar larvae to newly emerged adult stage. Thereafter, we tested their feeding response to five different types of food, i.e., *Aphis craccivora* Koch, *Aphis nerii* Boyer de Fonsclombe, conspecific eggs, heterospecific eggs and mixed pollen grains, equidistantly placed in Petri dish. On the basis of newly emerged adults’ food choice, they were reared on the same diet for 10 days. Thereafter, adults were paired with their opposite sex (collected from stock culture reared on *A*. *craccivora*) and mating and reproductive parameters were recorded. Our results suggested that the variety of food did not affect the preference of ladybird beetle, *P. dissecta*. They tend to choose their natural diet, *i.e*., aphid in each dietary regime. We found that previous dietary regime significantly influences the mating and reproductive parameters of both the male and female except for the time to commence mating by the male. Food choices of adult beetles were found to significantly influence the time to commence mating, average fecundity and percent egg viability in males and the mating duration in females.

## Introduction

Animals encounter many situations during their lifetime where they need to make a choice that may later prove crucial for their survival. For instance, which food to eat, whom to mate with and where to live etc. The consequences of poor decisions can be negative or even fatal. Therefore, animals should have evolved mechanisms that help them to make ‘good’ decisions (MacArthur and Pianka 1966; Rapoport 1989; Stephens and Krebs 1986). Decision-making, at its core, involves selecting among various alternatives, each of which can result in both favorable and unfavorable consequences. As a result, certain decision-making scenarios become intricate as individuals must meticulously weigh the pros and cons of each choice prior to making a wise decision.

In traditional models of animal and human choice, individuals aim to maximize their benefits from their choices. In terms of food selection, animals may be governed by basic consideration that effort for hunting can be done as long as the energy gained exceeds the energy lost (MacArthur and Pianka 1966). Based on this simple consideration, the optimal foraging theory (Pyke, 1984) can be used to understand the feeding behaviour of predators (Emlen 1966; MacArthur and Pianka 1966; Krebs et al. 1978). This theory generally takes the univariate approach to explain the decisions of individuals on the basis of intrinsic properties of food, including nutrient concentration and ampleness. Since the food environment is inherently diverse, foraging decisions are influenced by the interactions between multiple components of food and the forager. Individuals’ food choices are influenced by their past experiences in the biochemical contexts within which foods are consumed with nutrient contents (Villalba and Provenza 2009; Villalba et al. 2004; Forbes and Kyriazakis 1995). In addition, past food experiences have the potential to influence individual’s present food preferences and intake. Numerous studies in holometabolous insects have investigated the impact of larval diet on female selection of host plants. This phenomenon has sparked debate and scrutiny among various researchers (Emden et al. 1996, Barron 2001, Rietdorf and Steidle 2002, and Janz et al. 2009). Nevertheless, recent instances indicate that the experiences of larvae may affect the host selection behavior of adult insects in multiple species (Rietdorf and Steidle 2002, Akhtar and Isman 2003, Gandolfi et al. 2003, Hora et al. 2005, Olsson et al. 2006, and Moreau et al. 2008).

Generalist insect predators consume a variety of prey species throughout their lives (Schenk and Bacher 2002; Wilby et al. 2005; Nagai 1991). However, they don’t necessarily target every insect they encounter (Crocker and Whitecomb 1980). These predators often exhibit clear preferences, attacking some insect species while avoiding others (Richards 1982; Houck 1986; Digweed 1993). Predaceous ladybird beetles have a diverse diet, including aphids, scale insects, psyllids, white flies, and other soft-bodied arthropods (Hodek and Honek 1996). Their prey falls into four main categories: essential prey, which supports reproduction and larval development; alternative prey, providing energy for survival; rejected prey, featuring aposematic coloration and chemicals that deter predators; and toxic prey are potentially harmful to ladybird beetles (Majerus 2016; Hodek and Honek 2012). Several factors influence their prey preferences, including prey quantity, quality, other characteristics (e.g., morphology, mobility, defense), the predator’s learning ability, environmental conditions (temperature, photoperiod), and genetics. Ladybird beetles adjust their search behavior based on prey availability, shifting from extensive and intensive searching (Biesinger and Haefner 2005; Ferran and Dixon 2013). Understanding these factors is crucial for effective biocontrol programs and laboratory mass multiplication efforts.

Ecological and nutritional plasticity, high intrinsic rates of increase, short life cycle, presence of sexual dimorphism and most abundance in agroecosystem are the most probable reasons for the selection of *Propylea dissecta* (Mulsant) (Coleoptera: Coccinellidae) as experimental model for this study. It is predaceous in nature preferably consume aphids followed by whiteflies (Hodek 1996; Pervez and Omkar 2004; Omkar and Mishra 2005). Although *Propylea* species have a diverse diet, their prey range is not as broad (Pervez and Omkar, 2004) as that of other beetles like, *H. axyridis* (Pervez and Omkar 2006), *C. septempunctata* (Hodek 1996) and *Adalia bipunctata* Linnaeus (Omkar and Pervez 2005). This information highlights the feeding behavior and prey preferences of *Propylea* species in comparison to other related beetles. Despite its potential as a predator, the smaller size of *P. dissecta* prevents it from effectively preying on a vast range of organisms. In addition, *Propylea* is not as widely distributed as the above-mentioned ladybirds, so its prey range is not very extensive. There is a positive correlation between prey suitability and performance in *P. dissecta* in terms of oviposition, growth, development, reproduction, and demographic attributes (Pervez and Omkar 2004). Preferences for prey are primarily influenced by palatability, prey size, and seasonality. Laboratory colonies of ladybird beetles are often maintained on natural prey that are costly, ephemeral and labour intensive. Additionally, the migration of some aphids between plant species can produce local extinctions of aphid prey sources for ladybird beetles. Because of these constraints on aphidophagous ladybird beetles, alternative foods, such as conspecific eggs, heterospecific eggs and pollen grains, can fill an important dietary void. However, in presence of more food choices, beetle may end up confused and probability of choosing a poor food may increase. To test, if more food choices lead to bad results, we hypothesized that on being provided with variety of food choices, *P. dissecta* will show poor food selection, which in turn will negatively affect their mating and reproductive parameters.

## Material and Methodology

### Stock Culture

Adults of *Cheilomenes sexmaculata* Fabricius and *P. dissecta* were collected from the agricultural fields surrounding Lucknow, India (26°50’N 80°54’E). Adults were mated and placed in Petri dishes (hereafter, 9.0 × 2.0 cm unless mentioned) in temperature-controlled chambers maintained at 25±2°C, 65±5% RH, 14L:10D photoperiod. They were provided with *ad libitum* daily replenished supply of *Aphis craccivora* Koch (Hemiptera: Aphididae), infested on *Vigna unguiculata* L. reared in a polyhouse (25±2°C, 65±5% R.H.). The eggs laid were collected every 24 hours and incubated under the above abiotic conditions until hatching. After hatching, first instar larvae were removed using a fine camel hair paint brush and assigned individually to clean experimental Petri dishes and provided with *ad libitum* supply of aphids. In order to acclimatize to laboratory conditions, rearing was done for one more generation. Thereafter, the requisite stages were used for experimentation.

### Food collection for dietary choice

For food choice experiment, five different foods (Heterospecific eggs, conspecific eggs, mixed pollen grains, *Aphis nerii* Boyer de Fonsclombe, *Aphis craccivora* Koch) were provided. For the collection of conspecific and heterospecific eggs, pair of ten-day-old male and female (n=50 pairs) of both species, *i.e*. *P. dissecta* and *C. sexmaculata*, were taken from stock and allowed to mate in plastic Petri dishes under abiotic conditions mentioned above and provided with *ad libitum* prey. The females were isolated post mating in Petri dishes (biotic and abiotic factors as above) and observed for oviposition for five days. Fresh eggs laid were collected daily and only fresh eggs were given as food choice in the experiment. Aphids, *A. craccivora* and *A. nerii*, were collected from their host plants *V. unguiculata* and *Calotropis procera* (Aiton), respectively. Both the aphids were provided on the leaves of their host plants during food choice treatments to restrict their mobility. Bee collected mixed pollen grains used in the experiment were purchased from Royal Honey and Bee Farming Products (Nitinsbees), Chinhat, Lucknow, Uttar Pradesh.

### Optimization of food in each dietary regime

In order to evaluate the effect of dietary regime on multiple foods, dietary regimes were maintained. Standardization studies were first conducted to evaluate the food quantity to be provided in each regime and these food regimes were grouped as: (i) Aphid, *A. craccivora*, (ii) Aphid, *A. nerii*, (iii) freshly laid heterospecific eggs, (iv) freshly laid conspecific eggs, and (v) pollen grains. In aphid dietary regime the first, second and third instars of *P. dissecta* were provided with 6-12 second and third instars of *Aphid* per day and for the fourth instar,10-20 gravid aphids were provided. In egg dietary regime, the number of eggs provided differed with instars: first instar 20 eggs, second instar 40 eggs, third instar 60 and fourth instar with 100 eggs. For the pollen grains dietary regime, first crushed pollen grains were provided to the instars but only pollen fall short to maintain larval development. Then we provided the pollen grains along with cotton ball soaked with honey syrup. Here also we recorded the highest mortality of larvae. Out of 378 larvae only 49 survived on this diet. From these 49 we took 30 larvae for further experiment as pollen dietary regime. During whole study we referred pollen plus honey syrup diet as pollen grain dietary regime.

### Experimental Design

10-day-old adults of *P. dissecta* were taken from the stock culture and paired in plastic Petri dishes. After natural disengagement, males were removed and females were placed in Petri dishes individually. After every 24 hours, eggs laid were collected and incubated under the above-mentioned conditions until hatching.

### (1) Food choice trials

Immediately after hatching, first instars were placed individually in Petri dishes containing sufficient amount of food with the help of fine camel hair paint brush. Based on their food they were divided into following dietary groups for further rearing: (i) Aphid, *A. craccivora*, (ii) Aphid, *A. nerii*, (iii) freshly laid heterospecific eggs, (iv) freshly laid conspecific eggs, and (v) pollen grains. Optimum food in each dietary treatment were provided and replenished daily until pupation. Soon after emergence, adults were individually placed in large plastic Petri dish (14.5 cm diameter ×2.0 cm height) containing the food options, *i.e.*, (i) *A. craccivora*, (ii) *A. nerii*, (iii) Conspecific eggs, (iv) Heterospecific eggs, and (v) Pollen grains, placed equidistantly (Fig. 1)

**Figure 1.**
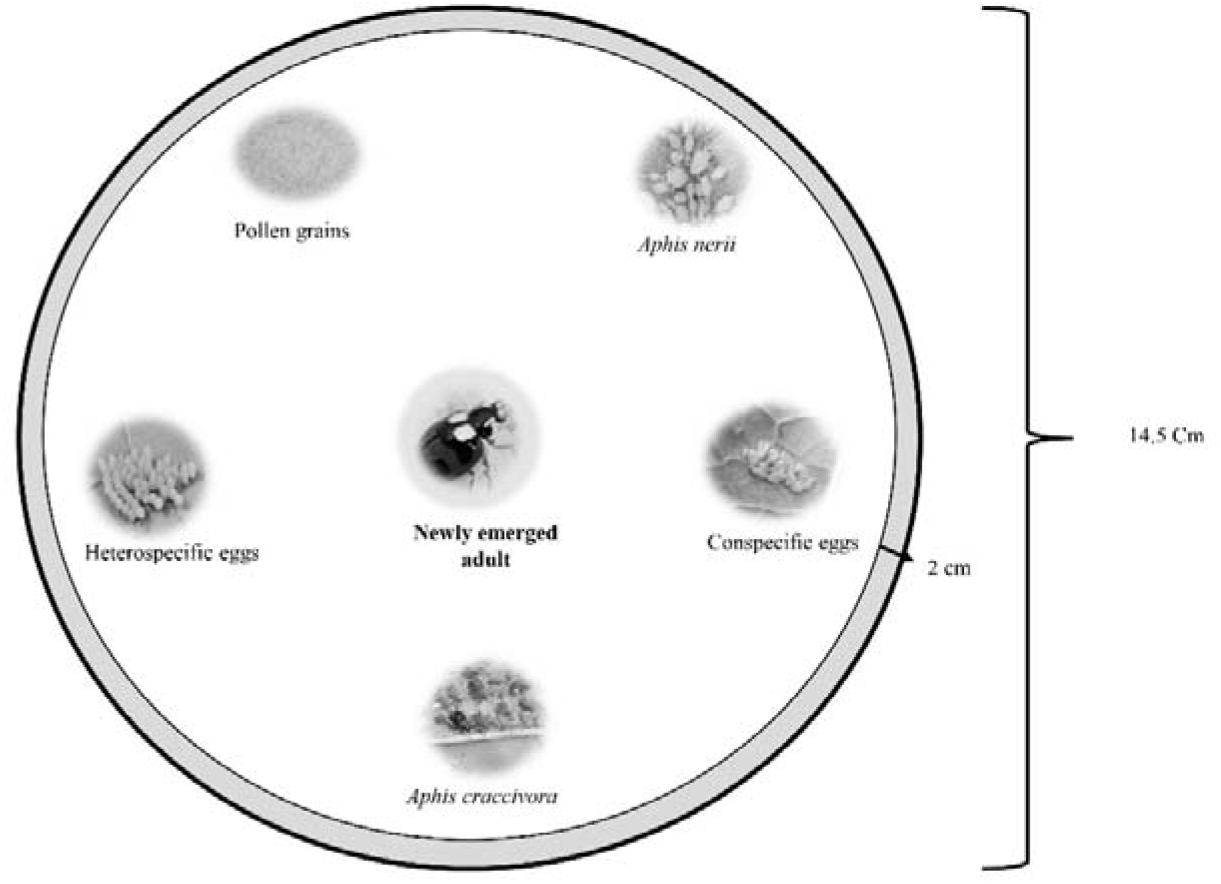
Diagrammatic representation of food choice trail setup.

Total developmental duration (from egg to adult emergence), first food encountered, time to encounter first food, first food choice, and time taken to consume first food were recorded. All the treatments were replicated 30 times.

### (2) Effect of preferred diet quality on mating and reproductive parameters

Newly emerged adults (either male or female) were reared on same diet on which they fed as larvae for the next 10 days. On reaching sexual maturity, adults were paired with their opposite gender of same age from stock culture. All the mating and reproductive parameters, such as, time to commence mating (TCM), copulation duration, preoviposition period, fecundity and percent egg viability was recorded. For the females’, we have checked the mating and reproductive parameter of its own but for the males we have analysed these parameters for females reared on *A. craccivora* taken from stock culture and paired with the males from different dietary regimes.

### Statistical analysis

Data on total development duration, time to encounter first food and time to consume first food were first tested for normality (Kolmogorov-Smirnoff test). On being found non-normally distributed, Generalised Linear Model (GLM) was applied. To analyse the effect of dietary regime on total development duration, the latter was used as response factor and former as fixed factor during GLM. To study the effect of multiple food choice on time to encounter first food, encounter time was taken as response factors and dietary regime and first food encountered were used as fixed factors in GLM. To reveal the effect of multiple food choice on time to consume first food, first consumption time was used as response factors and dietary regime and first food consumed were taken as fixed factors in GLM.

Chi-square (χ^2^) Goodness-of-fit analysis was used to analyse the first encountered food and first consumed prey (food choice) by the newly emerged adult. To test the effect of food choice on mating and reproductive parameters of adults, data on time to commence mating, copulation duration, oviposition period, average fecundity and percent egg viability were first tested for normality and on getting non normally distributed, GLM was applied. Data on time to commence mating, copulation duration, oviposition period, average fecundity and percent egg viability were taken as response factors and dietary regime and food choice were used as fixed factors in GLM.

All the analyses were carried out by using SPSS 20 software, version 20.0 (IBM, Armonk, New York, United States of America).

## Results

### (1) Effect of dietary regime on total development duration

The total developmental duration was significantly affected by the diet (F =781.271, df=4, P=0.000). Shortest development duration was recorded on *A. craccivora*, whereas longest duration was found on mixed pollen grains (Fig.2)

**Figure 2.**
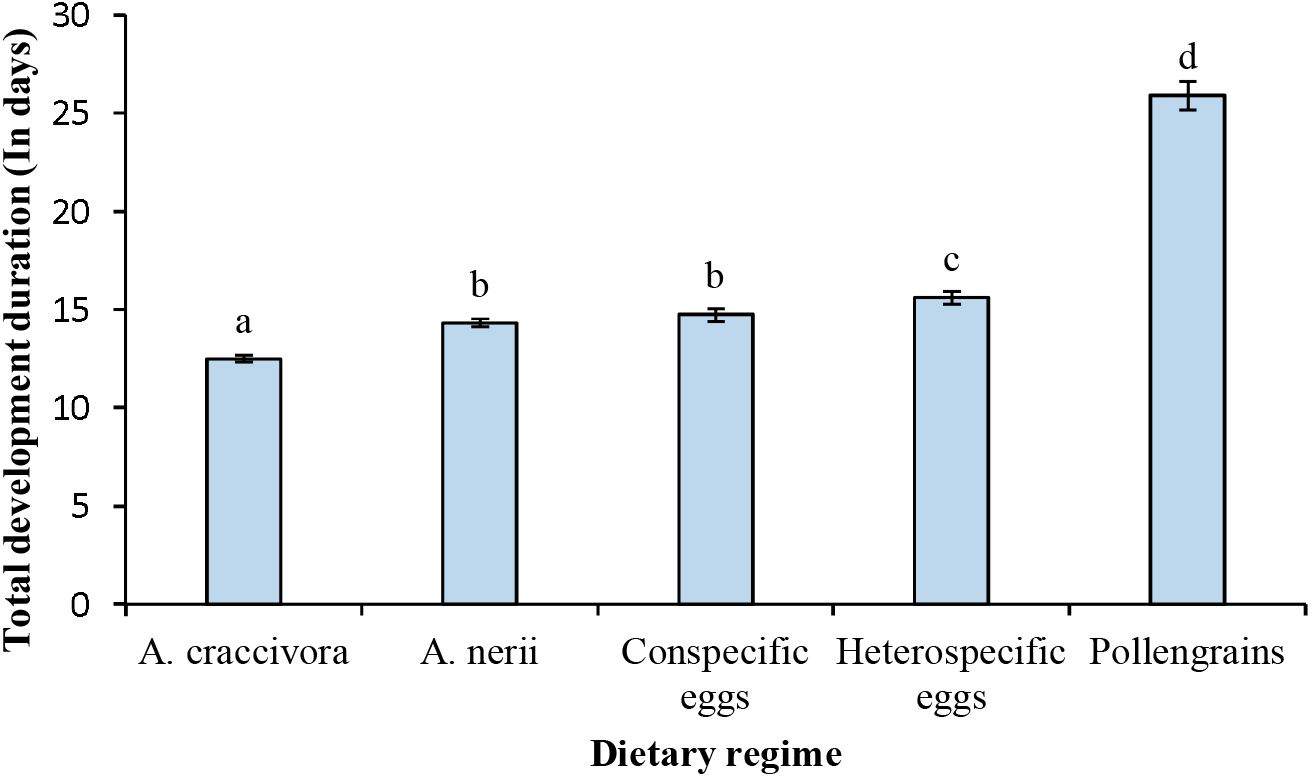
Values are mean ± SE. Different letters indicate comparison of means among different crowding regimes. Data to be significant (F = 781.271, df = 4, P = 0.000).

### (2) Food choice trials

Time to encounter first food was insignificantly affected by diet (F =2.523, df=4, P=0.640); irrespective of the type of food first encountered (F=4.150, df=4, P=0.386). The interaction between dietary regime and food first encountered was also insignificant (dietary regime*first food encountered: F =21.144, df=16, P=0.173) (Fig. 3). First encountered food was insignificantly (χ^2^ =21.446, df=16, P=0.162) affected by the dietary regime of the larvae. First encounters with food were random (Fig.4).

**Figure 3.**
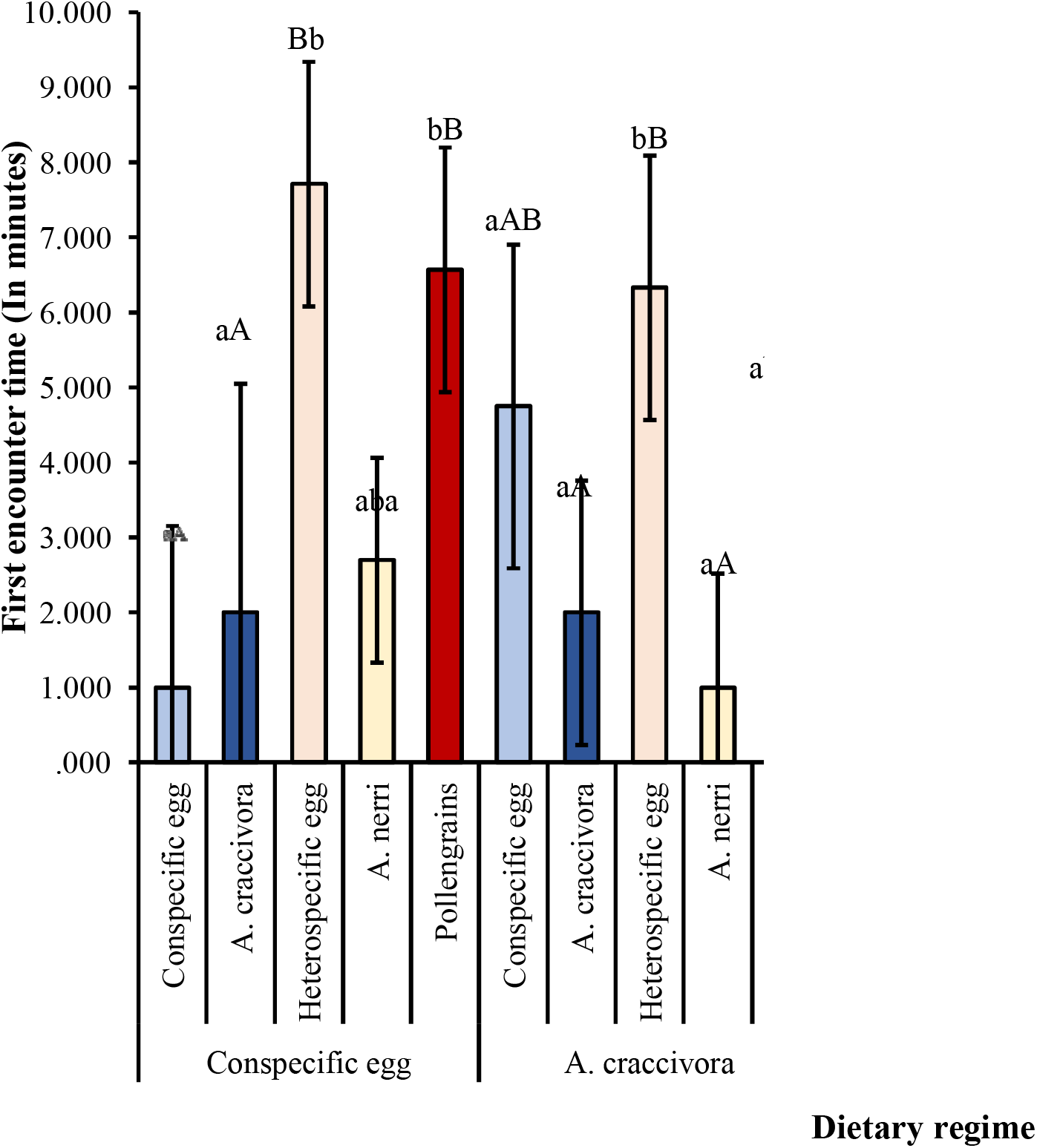
First encounter time (in seconds) in each dietary regime. Values are mean ± SE. Lowercase and uppercase letters indicate comparison of means within diet treatments and among different dietary regimes, respectively. Similar letters indicate lack of significance difference (P-value>0.05).

**Figure 4.**
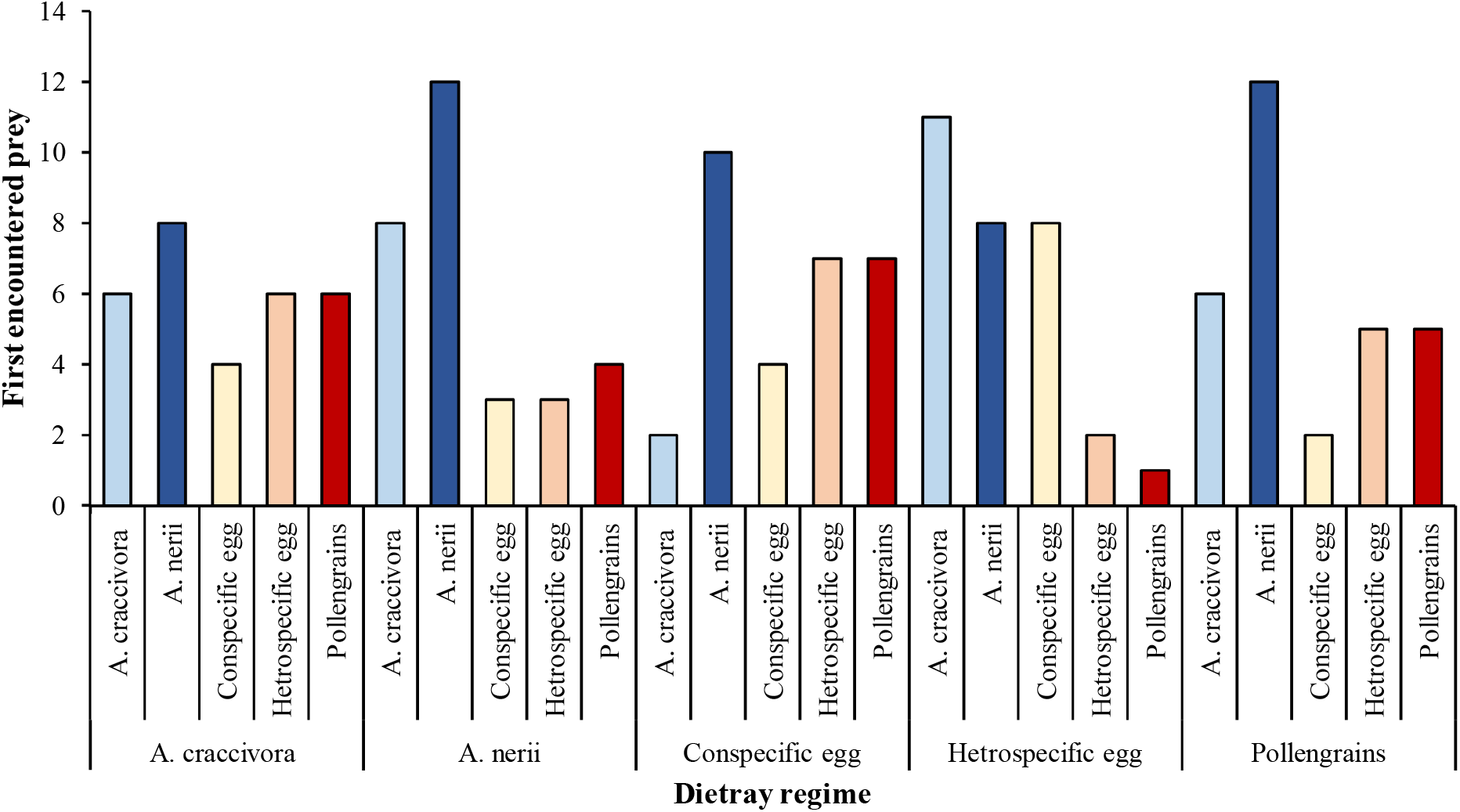
First food encountered in each dietary regime. Data are insignificant (χ^2^ =21.446, df=16, P=0.162).

Time to consume first food was insignificantly affected by both the dietary regime (F =5.476, df=4, P=0.242) and first food consumed (F =1.341, df=4, P=0.854). The interaction amongst two factors were insignificant (F =14.179, df=16, P=0.585). Similar consumption time was recorded in each dietary regime (Fig. 5). First consumed food was also found insignificant across dietary regimes (χ^2^ =15.671, df=16, P=0.476. In each dietary regime, first consumed food was either *A. craccivora* or *A*. *nerii* (Fig. 6)

**Figure 5.**
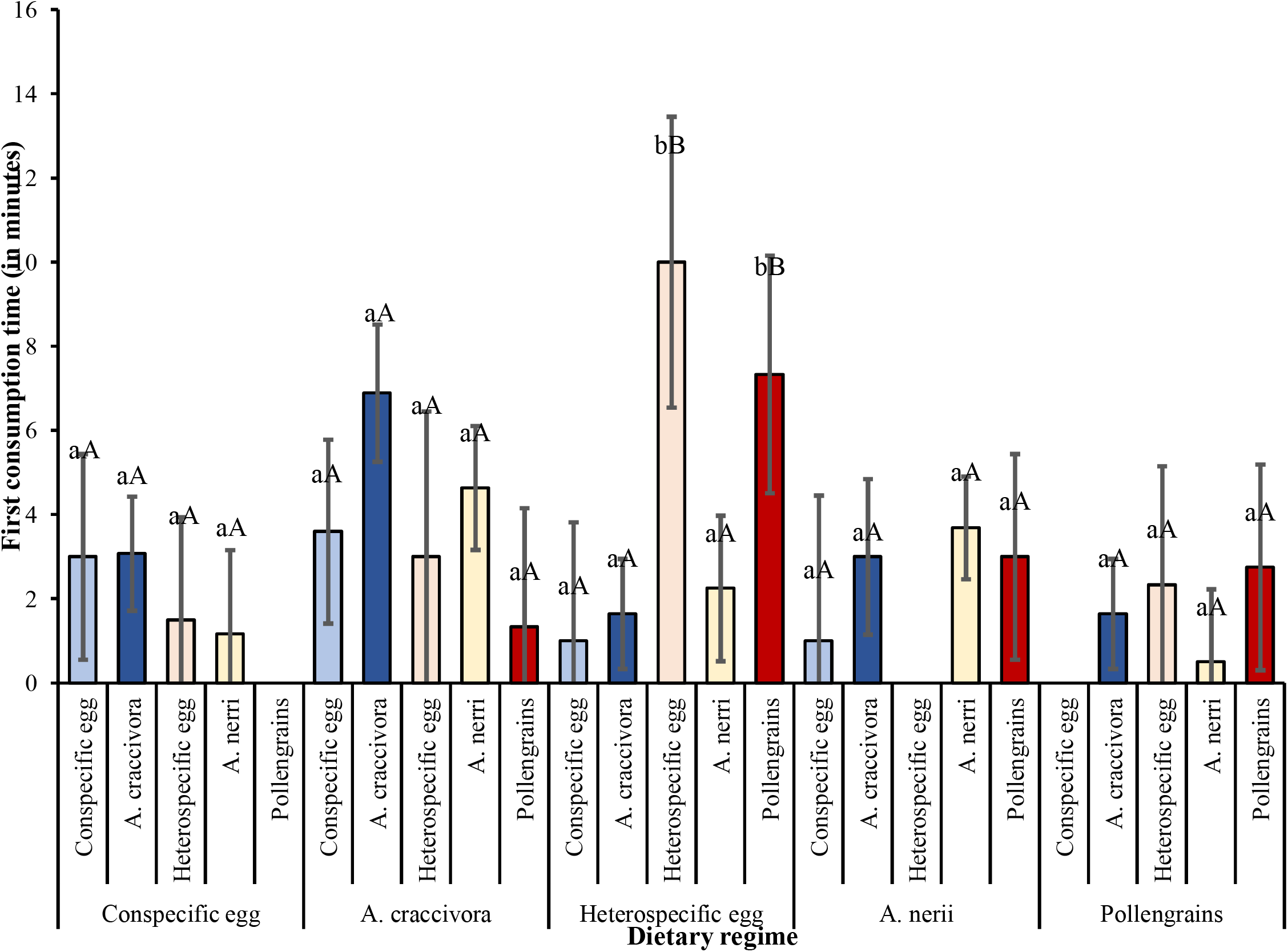
First consumption time in to consumed first food in each dietary regime. Values are mean ± SE. Lowercase and uppercase letters indicate comparison of means within diet treatments and among different dietary regimes, respectively. Similar letters indicate lack of significance difference (P-value>0.05).

**Figure 6.**
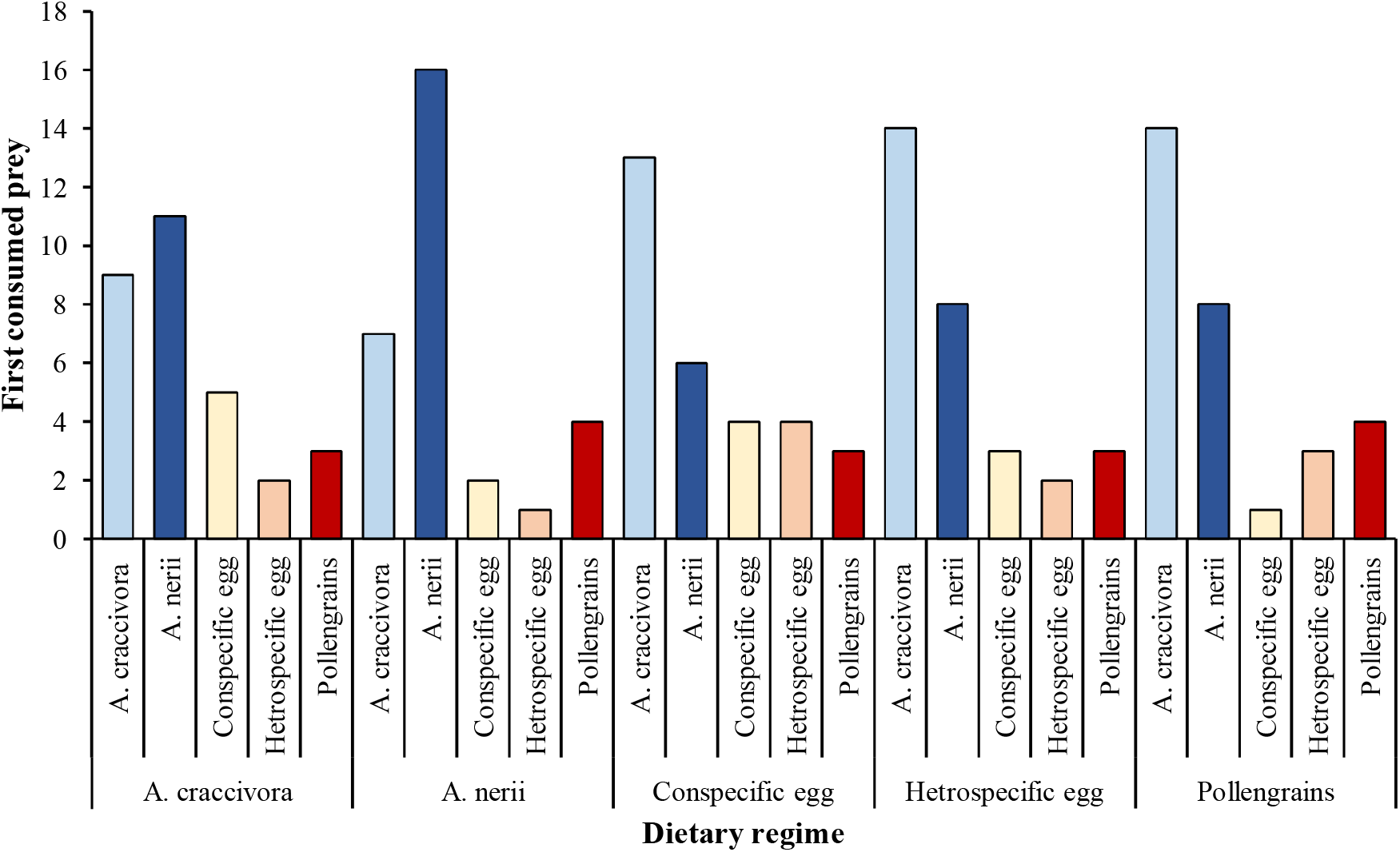
First consumed food in each dietary regime. Data are insignificant (χ^2^ =15.671, df=16, P=0.476).

### (3) Effect of selected diet quality on mating and reproductive parameters

Here, we assessed the effect of selected food on beetles’ mating and reproductive parameters. For male, TCM was significantly (F =14.363, df=4, P=0.006) affected by food choice they have made on day of their emergence but it was not influenced by their larval dietary regime (F =2.521, df=4, P=0.641). The interaction of first food choice and dietary regime was significant (F =62.729, df=14, P=0.000). Copulation duration was significantly (F =9.944, df=4, P=0.041) affected by dietary regime but not by food choice (F =5.128, df=4, P=0.274). However, the interaction between the two was significant (F =27.065, df=14, P=0.019) (Table 1)

**Table 1.**
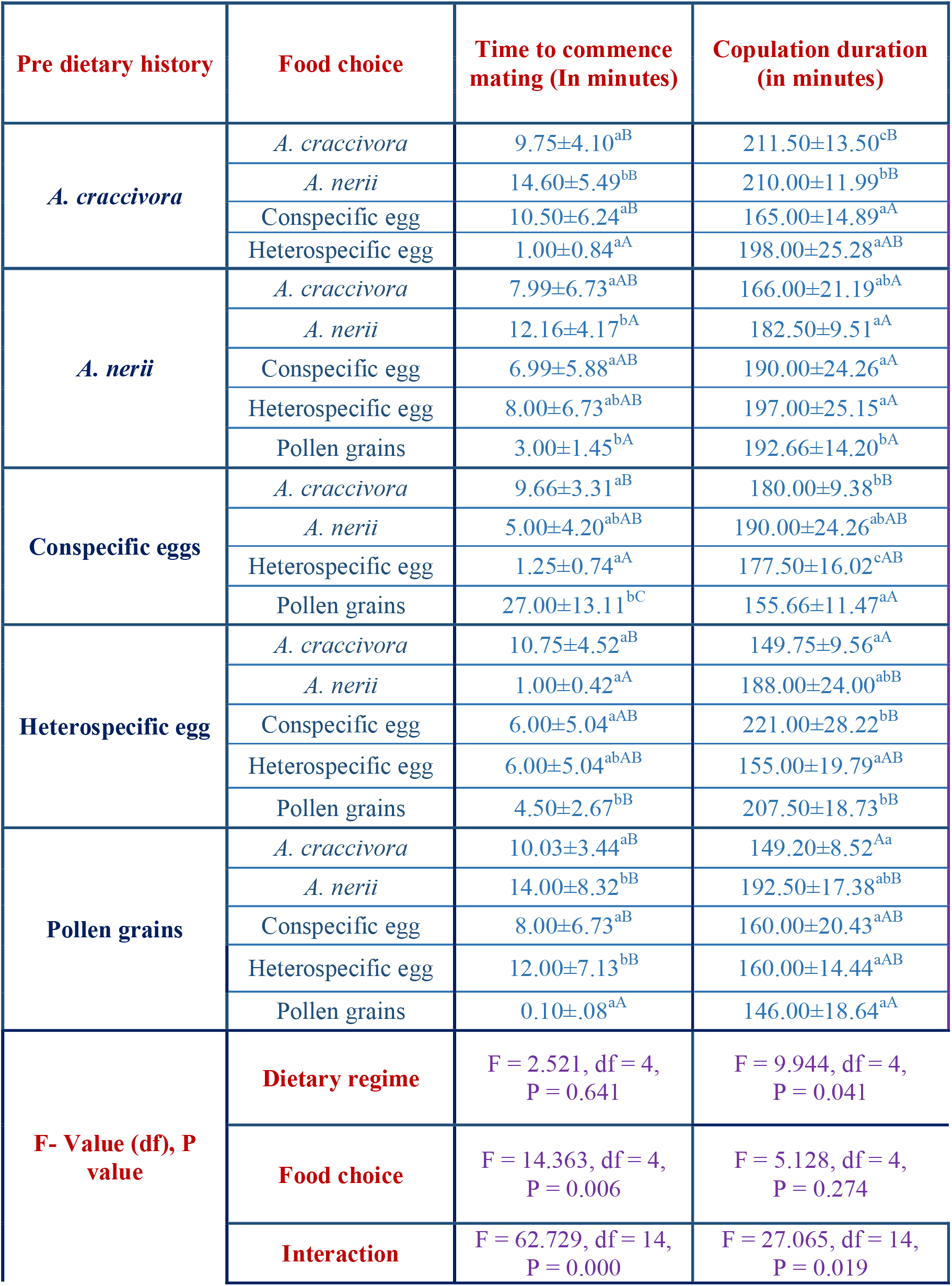
Effect of dietary regime and food choice on mating parameters of male *Propylea dissecta*. Values are mean ± SE. Letters indicate comparison of means among different dietary regimes, respectively. Means that do not share a letter are significantly different (P-value>0.05)

| Pre dietary history | Food choice | Time to commence mating (In minutes) | Copulation duration (in minutes) |
| --- | --- | --- | --- |
| *A. craccivora* | *A. craccivora* | 9.75±4.10<sup>aB</sup> | 211.50±13.50<sup>cB</sup> |
|  | *A. nerii* | 14.60±5.49<sup>bB</sup> | 210.00±11.99<sup>bB</sup> |
|  | Conspecific egg | 10.50±6.24<sup>aB</sup> | 165.00±14.89<sup>aA</sup> |
|  | Heterospecific egg | 1.00±0.84<sup>aA</sup> | 198.00±25.28<sup>aAB</sup> |
| *A. nerii* | *A. craccivora* | 7.99±6.73<sup>aAB</sup> | 166.00±21.19<sup>abA</sup> |
|  | *A. nerii* | 12.16±4.17<sup>bA</sup> | 182.50±9.51<sup>aA</sup> |
|  | Conspecific egg | 6.99±5.88<sup>aAB</sup> | 190.00±24.26<sup>aA</sup> |
|  | Heterospecific egg | 8.00±6.73<sup>abAB</sup> | 197.00±25.15<sup>aA</sup> |
|  | Pollen grains | 3.00±1.45<sup>bA</sup> | 192.66±14.20<sup>bA</sup> |
| Conspecific eggs | *A. craccivora* | 9.66±3.31<sup>aB</sup> | 180.00±9.38<sup>bB</sup> |
|  | *A. nerii* | 5.00±4.20<sup>abAB</sup> | 190.00±24.26<sup>abAB</sup> |
|  | Heterospecific egg | 1.25±0.74<sup>aA</sup> | 177.50±16.02<sup>cAB</sup> |
|  | Pollen grains | 27.00±13.11<sup>bC</sup> | 155.66±11.47<sup>aA</sup> |
| Heterospecific egg | *A. craccivora* | 10.75±4.52<sup>aB</sup> | 149.75±9.56<sup>aA</sup> |
|  | *A. nerii* | 1.00±0.42<sup>aA</sup> | 188.00±24.00<sup>abB</sup> |
|  | Conspecific egg | 6.00±5.04<sup>aAB</sup> | 221.00±28.22<sup>bB</sup> |
|  | Heterospecific egg | 6.00±5.04<sup>abAB</sup> | 155.00±19.79<sup>aAB</sup> |
|  | Pollen grains | 4.50±2.67<sup>bB</sup> | 207.50±18.73<sup>bB</sup> |
| Pollen grains | *A. craccivora* | 10.03±3.44<sup>aB</sup> | 149.20±8.52<sup>Aa</sup> |
|  | *A. nerii* | 14.00±8.32<sup>bB</sup> | 192.50±17.38<sup>abB</sup> |
|  | Conspecific egg | 8.00±6.73<sup>aB</sup> | 160.00±20.43<sup>aAB</sup> |
|  | Heterospecific egg | 12.00±7.13<sup>bB</sup> | 160.00±14.44<sup>aAB</sup> |
|  | Pollen grains | 0.10±.08<sup>aA</sup> | 146.00±18.64<sup>aA</sup> |
| F- Value (df), P value | Dietary regime | F = 2.521, df = 4,<br>P = 0.641 | F = 9.944, df = 4,<br>P = 0.041 |
|  | Food choice | F = 14.363, df = 4,<br>P = 0.006 | F = 5.128, df = 4,<br>P = 0.274 |
|  | Interaction | F = 62.729, df = 14,<br>P = 0.000 | F = 27.065, df = 14,<br>P = 0.019 |

Pre oviposition period was insignificantly (F=3.552, df=4, P=0.470) influenced by the food choice but significantly (F=15.625, df=4, P=0.004) influenced by the dietary regime. The interaction of food choice and dietary regime (F=42.327, df=14, P=0.000) was significant. Average fecundity was significantly affected by both dietary regimes (F=46.628, df=4, P=0.000) and food choice (F=15.882, df=4, P=0.003) and their interaction was also significant (F=60.022, df=14, P=0.000). Percent egg viability was also significant with respect to both the dietary regime (F=64.871, df=4, P=0.000) and food choice (F=16.966, df=4, P=0.002) and their interaction was also significant (F=85.864, df=14, P=0.000) (Table 2)

**Table 2.** Effect of dietary regime and food choice on reproductive parameters of male *Propylea dissecta*. Values are mean ± SE. Letters indicate comparison of means among different dietary regimes, respectively. Means that do not share a letter are significantly different (P-value>0.05)

| Pre dietary history | Food choice | Preoviposition period (in days) | Average fecundity | Percent egg viability |
| --- | --- | --- | --- | --- |
| *A. craccivora* | *A. craccivora* | 1.75±0.56<sup>aA</sup> | 20.55±2.88<sup>bA</sup> | 82.01±10.10<sup>bA</sup> |
|  | *A. nerii* | 2.80±0.50<sup>bA</sup> | 16.20±5.76<sup>aA</sup> | 96.70±20.21<sup>aA</sup> |
|  | Conspecific egg | 1.50±0.80<sup>aA</sup> | 18.00±2.35<sup>aA</sup> | 84.99±8.25<sup>aA</sup> |
|  | Heterospecific egg | 5.00±1.13<sup>bB</sup> | 18.00±2.88<sup>abA</sup> | 74.67±10.10<sup>bA</sup> |
| *A. nerii* | *A. craccivora* | 2.00±1.13<sup>aAB</sup> | 5.96±2.57<sup>Aa</sup> | 30.02±9.04<sup>aA</sup> |
|  | *A. nerii* | 2.16±0.46<sup>bB</sup> | 27.7±2.35<sup>bB</sup> | 93.84±8.25<sup>aB</sup> |
|  | Conspecific egg | 2.00±1.13<sup>abAB</sup> | - | - |
|  | Heterospecific egg | 3.00±1.13<sup>abB</sup> | 4.5±2.88<sup>aA</sup> | 22.44±10.10<sup>aA</sup> |
|  | Pollen grains | 0.99±0.65<sup>aA</sup> | 6.2±4.07<sup>aA</sup> | 39.88±14.29<sup>Aa</sup> |
| Conspecific eggs | *A. craccivora* | 2.00±0.46<sup>aA</sup> | 4.60±4.06<sup>aA</sup> | 37.73±14.298<sup>aA</sup> |
|  | *A. nerii* | - | 22.00±5.76<sup>abB</sup> | 93.24±20.21<sup>aB</sup> |
|  | Heterospecific egg | - | 22.6±5.76<sup>bB</sup> | 95.39±20.21<sup>bB</sup> |
|  | Pollen grains | - | - | - |
| Heterospecific egg | *A. craccivora* | 1.50±0.56<sup>aB</sup> | 16.00±5.76<sup>bAB</sup> | 97.49±20.21<sup>bB</sup> |
|  | *A. nerii* | 0.25±0.56<sup>aA</sup> | 23.4±5.76<sup>abB</sup> | 92.59±20.21<sup>aB</sup> |
|  | Conspecific egg | 4.00±1.13<sup>bC</sup> | - | - |
|  | Heterospecific egg | 1.00±1.13<sup>aAB</sup> | 10.6±5.76<sup>aA</sup> | 66.79±20.21<sup>bAB</sup> |
|  | Pollen grains | 1.50±0.80<sup>abAB</sup> | 8.6±4.07<sup>aA</sup> | 37.07±14.29<sup>aA</sup> |
| Pollen grains | *A. craccivora* | - | 25.00±9.11<sup>bA</sup> | 119.60±31.97<sup>bA</sup> |
|  | *A. nerii* | 2.00±0.80<sup>bA</sup> | 27.2±3.33<sup>bA</sup> | 90.73±11.67<sup>aA</sup> |
|  | Conspecific egg | - | - | - |
|  | Heterospecific egg | 3.00±0.80<sup>aA</sup> | 22.6±4.07<sup>bA</sup> | 78.81±14.29<sup>bA</sup> |
|  | Pollen grains | 3.00±1.13<sup>bA</sup> | 20.40±5.76<sup>bA</sup> | 81.86±20.21<sup>bA</sup> |
| F- Value (df), P value | Dietary regime | F = 15.625, df = 4, P = 0.004 | F = 46.628, df = 4, P = 0.000 | F = 64.871, df = 4, P = 0.000 |
|  | Food choice | F = 3.552, df = 4, P = 0.470 | F = 15.882, df = 4, P = 0.003 | F = 16.966, df = 4, P = 0.002 |
|  | Interaction | F = 42.327, df = 14, P = 0.000 | F = 60.022, df = 14, P = 0.000 | F = 85.864, df = 14, P = 0.000 |

For the female, TCM was insignificantly (F=2.129, df=4, P=0.712) affected by the food choice but was significantly (F =9.697, df=4, P=0.046) affected by the dietary regime. The interaction between food choice and dietary regime (Food choice*Dietary regime) was insignificant (F=8.600, df=10, P=0.570). Copulation duration was significantly influenced by both dietary regime (F =38.298, df=4, P=0.000) and food choice (F =10.962, df=4, P=0.027). Their interaction was also significant (F =50.980, df=10, P=0.000) (Table 3)

**Table 3.** Effect of dietary regime and food choice on mating parameters of female *Propylea dissecta*. Values are mean ± SE. Letters indicate comparison of means among different dietary regimes, respectively. Means that do not share a letter are significantly different (P-value>0.05)

| Pre dietary history | Food choice | Preoviposition period (in days) | Average fecundity | Percent egg viability |
| --- | --- | --- | --- | --- |
| Pre dietary history | Food choice | Time to commence mating (In minutes) | Copulation duration (in minutes) |  |
| *A. craccivora* | *A. craccivora* | 5.01±3.95<sup>aA</sup> | 180.00±19.55<sup>bA</sup> |  |
|  | *A. nerii* | 14.20±3.54<sup>abB</sup> | 201.60±17.49<sup>cB</sup> |  |
|  | Conspecific egg | 6.00±7.91<sup>abAB</sup> | 216.00±39.11<sup>bB</sup> |  |
|  | Pollen grains | 5.50±5.59<sup>aAB</sup> | 168.00±27.65<sup>abA</sup> |  |
| *A. nerii* | *A. craccivora* | 17.20±3.54<sup>bA</sup> | 151.20±17.49<sup>aA</sup> |  |
|  | *A. nerii* | 14.80±3.23<sup>bA</sup> | 189.00±15.96<sup>cB</sup> |  |
|  | Pollen grains | 7.00±7.91<sup>aA</sup> | 184.00±39.11<sup>bB</sup> |  |
| Conspecific eggs | *A. craccivora* | 5.40±3.54<sup>aA</sup> | 221.00±17.49<sup>cB</sup> |  |
|  | *A. nerii* | 13.33±4.57<sup>abAB</sup> | 196.33±22.58<sup>cB</sup> |  |
|  | Conspecific egg | 17.66±4.57<sup>bB</sup> | 158.00±22.58<sup>aA</sup> |  |
|  | Heterospecific egg | 13.00±7.91<sup>aAB</sup> | 240.00±39.11<sup>aC</sup> |  |
| Heterospecific egg | *A. craccivora* | 6.15±3.23<sup>aA</sup> | 138.83±15.96<sup>bB</sup> |  |
|  | *A. nerii* | 1.66±4.57<sup>aA</sup> | 60.33±22.58<sup>aA</sup> |  |
|  | Conspecific egg | 4.25±5.59<sup>aA</sup> | 230.50±27.65<sup>bC</sup> |  |
| Pollen grains | *A. craccivora* | 5.00±3.23<sup>aA</sup> | 171.66±15.96<sup>bB</sup> |  |
|  | *A. nerii* | 6.66±4.57<sup>aA</sup> | 150.00±22.58<sup>bA</sup> |  |
|  | Pollen grains | 5.00±5.59<sup>aA</sup> | 143.50±27.65<sup>aA</sup> |  |
| F- Value (df), P value | Dietray regime | F = 9.697, df = 4, P = 0.046 | F = 38.298, df = 4, P = 0.000 |  |
|  | Food choice | F = 2.129, df = 4, P = 0.712 | F = 10.962, df = 4, P = 0.027 |  |
|  | Interaction | F = 8.600, df = 10, P = 0.570 | F = 50.980, df = 10, P <0.05 |  |

Preoviposition period was insignificantly influenced by both dietary regime (F =7.088, df=4, P=0.131) and food choice (F =1.557, df=4, P=0.816). Their interaction was also insignificant (F =8.827, df=10, P=0.549). Average fecundity was significantly influenced by dietary regime (F =11.152, df=4, P=0.025) but it was insignificant with respect to food choice (F =3.008, df=4, P=0.556). Their interaction was also insignificant (F =11.983, df=10, P=0.286). The percent egg viability was insignificant (F =4.625, df=4, P=0.328) with respect to the food choice but it was significantly affected by the dietary regime (F =14.554, df=4, P=0.006). Their interaction was also insignificant (F =12.664, df=10, P=0.243) (Table 4)

**Table 4.**
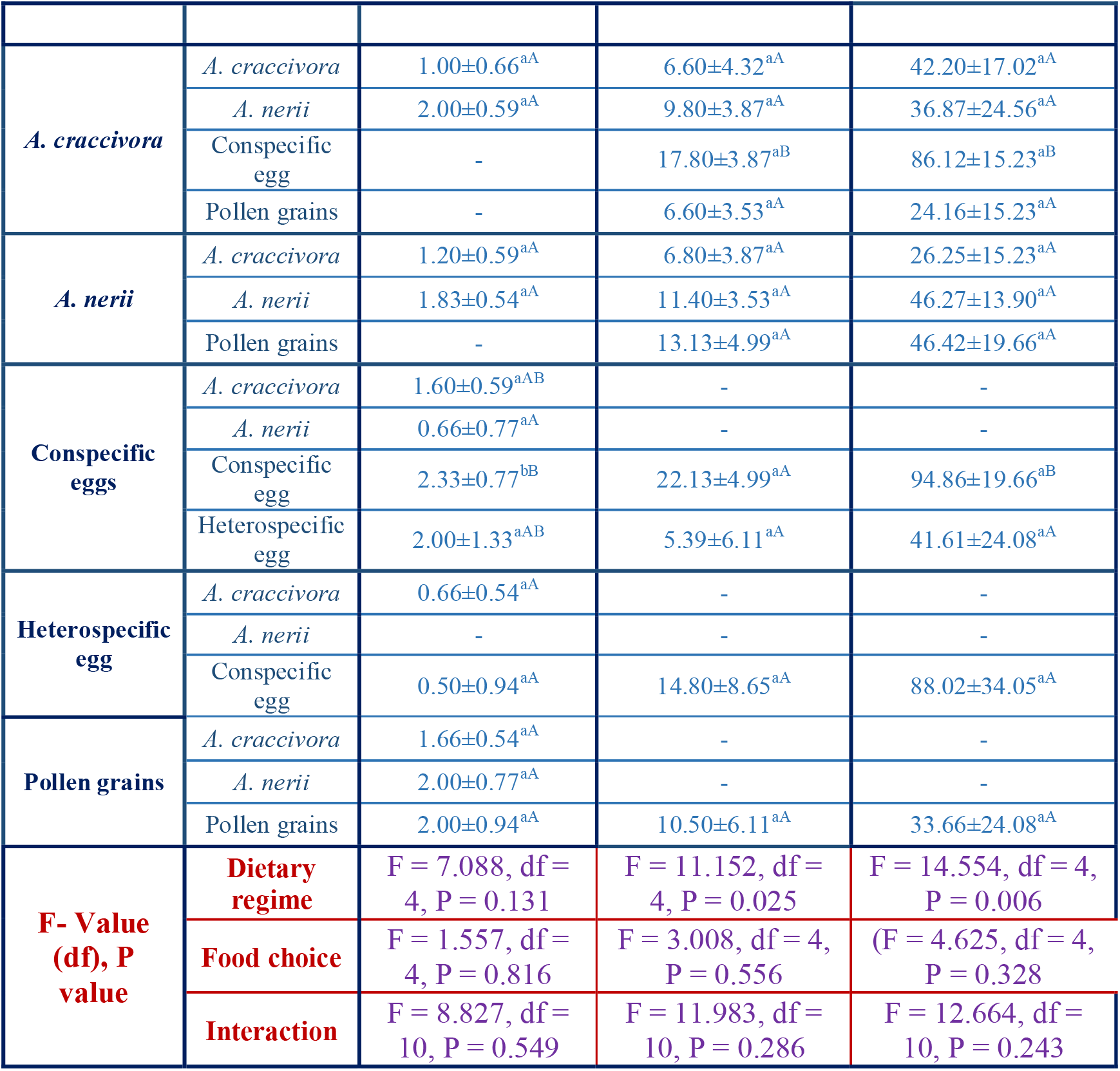
Effect of dietary regime and food choice on reproductive parameters of female *Propylea dissecta*. Values are mean ± SE. Letters indicate comparison of means among different dietary regimes, respectively. Means that do not share a letter are significantly different (P-value>0.05)

## Discussion

The present study was conducted to see the effect of too many food choices on food selection ability of the ladybird beetle and consequently affecting their reproductive physiology. However, our hypothesis was not supported and the preference for natural prey, in this case aphid was quite clear. The food choices made by males after emergence influenced time to commence mating, average fecundity and percent egg viability. The latter two factors along with copulation duration and pre-oviposition period were also influenced by dietary regime. Almost all recorded parameters were modulated by food choice and dietary regime.

The females’ previous dietary regime influenced all the reproductive and mating parameters whereas food choice they have made after emergence affected only the copulation duration. Significant interaction of food choice and dietary regime influenced the copulation duration and preoviposition period.

We used total developmental duration as an indicator of food suitability to *P. dissecta.* Decreasing developmental duration of *P. dissecta* is indicative of decreased food suitability, which in this case was A. *craccivora*> *A. nerii*> Conspecific eggs> Heterospecific eggs> Pollen grains. Earlier studies have indicated least suitability of *A. nerii* for *P. dissecta* (Pervez and Omkar 2004; Omkar and Mishra 2005). Osawa (2002) reported the faster development of immature stages of *Harmonia axyridis* (Pallas) on the diet of conspecific eggs as compared to non-cannibalistic diet. Similar result was reported in our finding as well where faster development of larvae on conspecific eggs over heterospecific eggs was recorded. Between the conspecific and heterospecific eggs, conspecific eggs were more nutritious to the larval stages than the heterospecific eggs (Kajita et al. 2010; Singh et al. 2020). Earlier it has already been established that *P. dissecta* slowed down its development on heterospecific eggs, possibly because they contain toxic alkaloids (Pervez et al. 2021). *P. dissecta* reared on pollen grains showed delayed development. Pollen grains may serve as an alternative source of food (Lima et al. 2020). In accordance with our findings, Farag et al. (2011) reported bee pollen as the least suitable diet for *Coccinella undecimpunctata* larvae, among seven different combinations of diets tested. Similar results were also reported for the larva of *Coccinella transversalis* Fab., where the larvae developed more efficiently when fed on honey and mealy bugs than on pollen or sugar syrup (Maurice et al. 2012).

First encounter time was random with all types of food. Even the first food encountered was found to be random. Similar findings were also reported for *Harmonia axyridis* where both adults and larvae searched randomly for prey-location (Canovai et al. 2019). This might be due to: (1) the arrangement of food, and (2) the type of food. On placing the newly emerged adult in middle of the arena surrounded by the variety of foods, enabled the beetle to first do extensive and then intensive search for food. Moreover, due to the presence of hemipteran and non-hemipteran food, *i.e.* conspecific eggs, heterospecific eggs and pollen grains, cues from these food items and the presence of sensory abilities in predators helped them to do extensive and then intensive search for all the food items. Earlier similar findings were reported that on being locating their prey, beetles tend to do intensive search (Ettifouri and Ferran 1993; YunDing et al. 1997). Being a generalist predator, *P. dissecta* not only move towards hemipterans but also towards non-hemipteran food types. It was reported that in dense prey colony beetles walk leisurely (Murdie 1971) and Hodek and Evan (2012) pointed them as “blundering idiots”.

Time to consume first food and first food consumed both were found to be unaffected by the previous dietary regimes. In each dietary treatment, adults tend to prefer their natural food over non hemipteran diets. This indicated the choosiness of newly emerged adult *P. dissecta*. Even on being provided with variety of food choices, they were found to be rigid in their food selection. Aphids being general nursery prey for aphidophagous ladybird beetle, *P. dissecta*, were preferred the most. On the other hand, pollen grains, conspecific and heterospecific eggs were least preferred in each dietary regime possibly due to the presence of less nutritive contents (Farag et al. 2011), insufficient immature stage of nutrition and toxic allelochemicals (Pervez et al. 2021), respectively. Overall comparison in different dietary regimes indicated food choice for aphids only. Thus, it indicates that the presence of multiple food varieties did not lead to poor food choice in *P. dissecta* and irrespective of the choices offered, beetles preferred their natural prey.

In males, time to commence mating was significantly influenced by their post emergence food choice rather than their overall dietary regime. The prolong time to commence mating in males with different food choices was indicative of them being not preferred by females reared on *A. craccivora*. This also suggested that females were able to identify the males on the basis of their dietary history. For instance, male crickets that have low nutritional concentrations or are provided with low-quality food are perceived as less attractive mates by females (Wilder and Rypstra 2007). In *Cryptolaemus montrouzieri* (Mulsant) female beetles exhibited a preference for mating with males that were raised with abundant nutritional diet, regardless of the nutritional conditions they themselves encountered during their larval development (Xie et al. 2015).

In females, we recorded opposite results. The time to commence mating was not impacted by the food choice but was affected by the dietary regime. Males reared on *A. craccivora* and females reared on *A. nerii* showed delayed time to commence mating than females of other dietary regimes. It indicated that poor dietary history of females changed the females’ attractiveness as mate. Similar to our findings in butterfly and other holometabolous insects, reproduction depends on the reserves accumulated during the larval dietary history (Bauerfeind and Fischer 2005).

Postemergence food choice by the adults did not affect males’ copulation duration but it did in females as males were consisted of females with different food choices and male reared on *A. craccivora*. This is indicative of the dependency of female copulation duration on food choice. In recent past it was observed that mating, being an energy consuming process, alters the food choices in females of *P. dissecta* (Verma et al. 2023). Interestingly, the copulation duration in both the sexes were significantly influenced by the dietary regime. Highest copulation duration was recorded for adults with larval dietary regime of *A. craccivora* followed by *A. nerii*, and conspecific eggs, whereas it was shortest in adults with larval dietary regime of heterospecific eggs and pollen grains. The lower copulation durations on larvae reared on heterospecific eggs and pollen grains can be explained with the fact that they were earlier considered to be toxic (Omkar and Pervez, 2011) or insufficient nutrients (Maurice et al. 2012).

Preoviposition period, a crucial phase in the reproductive cycle, remained unaffected by the food choice by adult insects but was notably impacted by the dietary regime of the individuals. This finding suggests that the availability of food sources at the time of adult eclosion does not play a significant role in determining the pre-oviposition period. This finding was consistent with previous study, it was found that the diet consumed at the moment of eclosion had no discernible effect on factors, such as preoviposition period, daily egg oviposition rates, or the viability of egg batches (Hatt and Osawa 2021). Findings underscore the importance of the dietary regime during the larval and developmental stages as a critical determinant of the preoviposition period with adult food choices having limited (Dmitriew and Rowe 2011).

In our study, we found that both the dietary regime and specific food choices played a significant role in determining the average fecundity and the percent egg viability in *A. craccivora* reared females when they were mated with males exhibiting different food choices. There might be two possible reasons for this result, (1) when females reared on preferred diet and males with the different food choices were mated, females enhanced their investment in the offspring. Similar to our findings, *Callosobruchus maculatus* (Fabricius) females laid more eggs in better food condition (Iglesias-Carrasco et al. 2018). With different food choices or diet switching from larval to adult stage might lead to male beetles poorly fed which resulted into enhanced investment of females in fecundity. In consistence with our results, swordtail females also enhanced their investment in offspring size and numbers when mated to poorly fed males (Kindsvater et al. 2013). (2) The other possible reason would be that male feeding treatment had no effect on female fecundity (Boggs and Freeman 2005).

Interestingly, it was found that the dietary history traits of females had a substantial impact on their average fecundity, while the food choices of females did not have a significant influence on their reproductive parameters. The possible reason for this finding could be that females having different food choices might be food deprived female perhaps because of food switching from larval to adult stage. This trend was also mirrored in the percentage of egg viability. These observations may be attributed to changes in the dietary regimes of both males and females during their emergence into adulthood, indicating the intricate interplay between food choice, dietary history, and the reproductive outcomes of these insects. In decorated crickets, *Gryllodes sigillatus* food-limited females, but not males, have been shown to decrease and delay their investment in reproduction (Houslay et al. 2015). Insignificant effect of females’ food choice and significant effect of males’ food choice on percent egg viability was recorded, it could be the nutritional needs of male *P. dissecta*, with regard to reproductive viability are different from the needs of females *P. dissecta*. Similar to our results, in *Podisus maculiventris* (Say) male nutritional demands are different from the females for the reproductive viability (Wittmeyer et al. 2010).

In conclusion, *P. dissecta* did not make bad decisions while interacting with different food regimes. They tend to select their natural and preferred food. However, dietary history of adult *P. dissecta* influenced the several life history parameters and their adult dietary regimes influenced their mating and reproductive parameters. Overall effect of the food choice was recorded in the males’ time to commence mating, average fecundity and precent egg viability but in females only copulation duration was affected by its own food choice.

